# Molecular understanding of *Eubacterium limosum* chemostat methanol metabolism

**DOI:** 10.1101/2022.11.04.514945

**Authors:** Jamin C. Wood, R. Axayacatl Gonzalez-Garcia, Dara Daygon, Gert Talbo, Manuel R. Plan, Esteban Marcellin, Bernardino Virdis

## Abstract

Methanol is a promising renewable energy carrier that can be used as a favourable substrate for biotechnology, due to its high energy efficiency conversion and ease of integration within existing infrastructure. Some acetogenic bacteria have the native ability to utilise methanol, along with other C_1_ substrates such as CO_2_ and formate, to produce valuable chemicals. Continuous cultures favour economically viable bioprocesses, however, the performance of acetogens has not been investigated at the molecular level when grown on methanol. Here we present steady-state chemostat quantification of the metabolism of *Eubacterium limosum*, finding maximum methanol uptake rates up to 640±22 mmol/gDCW/d, with significant fluxes to butyrate. To better understand metabolism of acetogens under methanol growth conditions, we sampled chemostats for proteomics and metabolomics. Changes in protein expression and intracellular metabolomics highlighted key aspects of methanol metabolism, and highlighted bottleneck conditions preventing formation of the more valuable product, butanol. Interestingly, a small amount of formate in methylotrophic metabolism triggered a cellular state known in other acetogens to correlate with solventogenesis. Unfortunately, this was prevented by post-translation effects including an oxidised NAD pool. There remains uncertainty around ferredoxin balance at the methylene-tetrahydrofolate reductase (MTHFR) and at the Rnf level.

## 1 Introduction

Acetogens are promising cellular factories for sustainable chemical and fuel production, due to their ability to close the carbon cycle using renewable carbon and electricity.^1^ These can be provided to cells through intermediates like hydrogen (H_2_), carbon monoxide (CO), formate and methanol. Methanol is already a widely used intermediate in the chemical industry, and can be produced renewably from carbon dioxide (CO_2_) for as little as US$560/t, which is expected to reach parity with fossil fuel derived methanol by 2032.^2^ Unlike utilisation of synthesis gas (syngas, a mixture of CO and H_2_), that has been commercialised by LanzaTech from waste,^3^ much less is understood about liquid C_1_ feedstocks such as methanol.^4^

Being liquid, methanol avoids transportation problems presented with gaseous substrates. For example, most chemical supply chains are distributed across the globe, and require long-distance transportation. That would likely be replicated for H_2_ used in syngas fermentations, especially if it was generated electrochemically in regions with low renewable electricity costs. Given that hydrogen transport results in non-negligible losses, this would add to costs and global warming due to hydrogen’s updated GWP of 11.^5^ On the other hand, methanol is more compatible with existing fermentation and supply chain infrastructure. As a liquid, methanol can also overcome key mass transfer limitations of gas fermentations and with higher energy efficiency, resulting in lower operational costs for things like cooling.^6^ Some acetogens, such as *Eubacterium limosum, Eubacterium callendari, Acetobacterium woodii*, and *Butyribacterium methylotrophicum*, have the unique ability to consume methanol. Exploring this capability further may overcome the need to engineer conventional model organisms.^7–9^

In acetogens, methanol enters the Wood-Ljungdahl Pathway (WLP) as methyl-THF via a methyltransferase,^10^ where mtaB catalyses the transfer of the methyl group to mtaC and is then transferred to the THF carrier catalysed by mtaA.^11,12^ It then results in partial oxidation (*i.e*., reversal of the WLP methyl branch) to generate reducing equivalents and satisfy CO_2_ demand for the carbonyl branch.^13,14^ However, sustained growth has not been shown with methanol as a solesubstrate, and a more oxidised co-substrate, such as CO_2_ or formate, is required.^15^ Yet, no physiological reason has been identified as to why this may be the case. Previously we have shown for *E. limosum*, that increasing the amount of methanol relative to the co-substrate resulted in more reduced products, such as butyrate, butanol and hexanoate. However, when the co-substrate was formate, distinct growth phases indicated balancing of metabolisms.^4^ Given that formate as the co-substrate had a significant growth rate advantage over CO_2_, likely because CO_2_ uptake rate is known to be a limitation,^16^ being able to understand and control co-consumption would allow better product specificity.

Molecular quantification and metabolic modelling have been used to improve understanding of metabolism, allowing for optimisation of process economics and fermentation designs.^17,18^ Whilst transcriptome^19^ and translatome^20^ data exist for *E. limosum*, these data sets were collected during batch fermentations for differential analysis of autotrophic and heterotrophic growth. Similarly, proteomic data of autotrophic, methylotophic and heterotrophic growth has been collected for closely related *E. callanderi*.^10^ However, acetogen metabolism is known to be controlled posttranscription and translation,^17,18,21^ and so metabolomic data may be vital to reveal growth bottlenecks. One research group conducted central-carbon metabolomic analysis for *E. limosum* methylotrophic growth in batch during the 1990’s,^15^ however, there have been significant advances in analytical methods since then. In contrast to these batch datasets, steady-state chemostat cultures have greater operational control and are more comparable between experiments, offering greater insight into process optimisation.^17^

Here, to understand methylotrophic growth and its limitations, we investigated steady-state C_1_ liquid methylotrophic growth in chemostat for *Eubacterium limosum. We* hypothesised that despite formate and CO_2_ being intermediates in the WLP, when they are co-substrates in methylotrophic fermentation, there are bottlenecks preventing assimilation at ratios supporting high product specificity. To test this, we undertook differential analysis between growth conditions, finding the differences are more related to thermodynamics than protein expression. We understand that a metabolic model is under construction for *E. limosum*,^22^ and we believe that the data presented here may help validate methylotrophic features.

## 2 Experimental

### 2.1 Bacterial Strain, Growth Medium, and Continuous Culture Conditions

*Eubacterium limosum* ATCC 8486 (*E. limosum)* was isolated as a single colony for all experiments after more than 200 generations of growth on 500 mM methanol and 100 mM formate. This phenotype was cultivated anaerobically in chemically defined phosphate buffered medium at 37 °C. Conditions are listed in **Table 1**. The medium comprised, per litre: 0.5 g MgCl_2_.6H_2_O, 0.5 g NaCl, 0.13 g CaCl_2_.2H_2_O, 0.75 g NaH_2_PO_4_, 2.05 g Na_2_HPO_4_, 0.25 g KH_2_PO_4_, 0.5 g KCl, 2.5 NH_4_Cl, 0.017 g FeCl_3_.6H_2_O, 0.5 g L-cysteine hydrochloride monohydrate, 1 mL of 1 g/L resazurin, 10 mL trace metal solution (TMS), 10 mL B-vitamin solution. The TMS comprised, per litre: 1.5 g nitrilotriacetic acid, 3 g MgSO_4_.7H_2_O, 0.5 g MnSO_4_.H_2_O, 1 g NaCl, 0.667 g FeSO_4_.7H_2_O, 0.2 g CoCl_2_.6H_2_O, 0.2 g ZnSO_4_.7H_2_O, 0.02 g CuCl_2_.2H_2_O, 0.014 g Al_2_(SO_4_)_3_.18H_2_O, 0.3 g H_3_BO_3_, 0.03 g Na_2_MoO_4_.2H_2_O, 0.028 g Na_2_SeO_3_.5H_2_O, 0.02 g NiCl_2_.6H_2_O and 0.02 g Na_2_WO_4_.2H_2_O. The B-vitamin solution comprised, per litre: 20 mg biotin, 20 mg folic acid, 10 mg pyridoxine hydrochloride, 50 mg thiamine-HCl, 50 mg riboflavin, 50 mg nicotinic acid, 50 mg calcium pantothenate, 50 mg vitamin B_12_, 50 mg 4-aminobenzoic acid and 50 mg thioctic acid. As the *E. limosum* Rnf complex is thought to be Na^+^ dependant, NaCl was supplemented to ensure all conditions had a consistent amount of sodium (*i.e*. as formate-containing chemostats used sodium formate as a substrate).

**Table 1.** Conditions summary of fermentations. N = stirrer speed; BR = biological replicates; pH = pH setpoint, BC = biomass concentration. Formate/0.4D is provided for comparison from ^23^.

| Identifier | Concentration (mM) |  | Gas | N | BR | D | pH | BC | Acetate | Butyrate |
| --- | --- | --- | --- | --- | --- | --- | --- | --- | --- | --- |
|  | Methanol | Formate |  |  |  |  |  |  |  |  |
|  |  |  |  | rpm | # | day<sup>-1</sup> |  | gDCW/L | g/L | g/L |
| Formate/0.4D | 0 | 100 | 100% Ar | 400 | 2 | 0.4 | 6.8 | 0.07±0.02 | 0.014±0.006 | 0.006±0.001 |
| MeOH/CO<sub>2</sub>/0.4D | 100 | 0 | 85% Ar,<br>15% CO<sub>2</sub> | 400 | 2 | 0.4 | 6.8 | 0.54±0.05 | 1.60±0.08 | 0.99±0.05 |
| 3MeOH/Formate/CO<sub>2</sub>/0.4D | 75 | 25 | 85% Ar,<br>15% CO<sub>2</sub> | 400 | 2 | 0.4 | 6.8 | 0.15±0.04 | 1.74±0.17 | 0.60±0.13 |
| 10MeOH/Formate/CO<sub>2</sub>/1D | 250 | 25 | 85% Ar,<br>15% CO<sub>2</sub> | 400 | 2 | 1 | 7.3 | 0.20±0.04 | 0.27±0.13 | 1.18±0.08 |
| 10MeOH/Formate/1D | 250 | 25 | 100% Ar | 400 | 2 | 1 | 7.3 | 0.10±0.01 | 0.19±0.02 | 0.60±0.18 |
| 10MeOH/Formate/2D | 250 | 25 | 100% Ar | 400 | 1 | 2 | 7.3 | 0.13±0.04 | 0.18±0.08 | 0.633±0.10 |

0.5 L Multifors bioreactors (infors AG) controlled by EVE software were operated at a working volume of 350 mL (magnetic marine impeller agitation), equipped with mass flow controllers (MFCs), peristaltic pumps, pH and temperature sensors. On-line off-gas analysis was conducted by connecting the system to a Hiden HPR-20-QIC mass spectrometer (Hiden Analytical). 10 μL/h antifoam (Sigma 435503) was added to the bioreactor. Steady-state results were obtained after gas uptake/production rates, optical density, and acid/base addition rates were constant for at least five working volumes. A second set of experiments were performed in 1 L Biostat B bioreactors (Sartorius) at a working volume of 500 mL (Rushton turbine agitation), also equipped with mass flow controllers (MFCs), peristaltic pumps, pH and temperature sensors, antifoam control, but connected to a Thermostar GSD 300T mass spectrometer (Balzers). Three technical replicate samples were collected per biological replicate, spaced by one dilution volume.

### 2.2 Experimental analysis and Quantification

#### 2.2.1 Extracellular analysis

Optical density (OD) was measured with a UV-Vis spectrophotometer (Thermo Fischer Scientific Genesys 10S UV-Vis Spectrophotometer, USA) at 600 nm. 0.32 gDCW/L/OD and a biomass formula of C_4_H_7_O_2_N_0.6_ was used to calculate molar cell concentrations.^4^ Extracellular metabolomic analysis liquid samples were collected, filtered and stored at −20 °C, before analysing by high performance liquid chromatography using an Agilent 1200 HPLC System with Phenomenex Rezex RHM-Monosaccharide H+ column (7.8 x 300 mm, PN: OOH-0132-KO) and guard column (Phenomenex SecurityGuard Carbo-H, PN: AJO-4490). 30 μL of sample was injected and analytes eluted isocratically with 4 mM H_2_SO_4_ at 0.6 mL min^-1^ for 48 min and a column oven temperature of 65 °C. Analytes were monitored using a UV detector (210 nm) and RID at positive polarity and 40 °C.

An on-line mass spectrometer, monitoring the amounts of H_2_, Ar and CO_2_ at 2, 40 and 44 amu respectively was used to analyse bioreactor off-gas. ‘On-line’ calibration was utilised, with a bypass line from the feed gas bottle assessed during each MS-cycle. Specific rates (mmol/gDCW/d) were calculated using bioreactor liquid working volume, pH and CO_2_ dissolution equilibrium, steady-state biomass concentration, ideal gas molar volume and the change in gas composition and flow rate (based on constant flow of inert Argon).

As a quality check, gaseous samples were also analysed ‘off-line’ with a Shimadzu 2014 GC. The system was equipped with a ShinCarbon packed column (ST 80/100, 2mm ID, 1/8 OD Silico, Restek), and H_2_ and Argon were monitored by a thermal conductivity detector (TCD), while CO_2_ was monitored using a flame ionization detector (FID).

A special note is made for methanol concentration and consumption calculations, given its relatively high volatility. Methanol losses from the feed and reactor were estimated using Henry’s law and then subtracted from consumption rates.

#### 2.2.2 Intracellular analysis

Direct data-independent acquisition mass spectrometry (direct-DIA) was used to carry out quantitative proteome analysis. Sample preparation was based on a method developed previously for autotrophic growth of *C. autoethangenum*, briefly discussed here. 5 ODs of culture was immediately pelleted by centrifugation (16,000g for 3 minutes at 4 °C), then washed with 1 mL PBS (Sigma P4417). A further centrifugation allowed the supernatant to be discarded. The remaining sample was stored at −20 °C for later analysis. Approximately 100 μg of protein was resuspended in 25 μL Milli-Q water and then combined with 25 μL SDS lysis buffer (10% SDS, 100 mM Tris). Formate grown cells required bead beating (0.1 mm diameter glass beads) to assist cell lyses. Dithiothreitol (DTT) was added to a final concentration of 20 mM, incubating at 70 °C for 1 hour to reduce protein disulphide bonds. To alkylate cysteine, Iodoacetamide (IAA) was added to a final concentration of 40 mM, incubating in the dark for 30 minutes. The reaction was then sonicated before adding 2.5 μL 12% phosphoric acid, and combining with 165 μL S-Trap binding buffer (90% methanol, 100 mM Tris (aq)). The sample was centrifuged at 13,000g for 8 minutes, and then added to the S-Trap spin column. It was spun for 1 minute at 4000g and thrice washed with 150 μL S-Trap binding buffer. 50 μL of 50 mM ammonium bicarbonate pH 8 with 1 μg of trypsin was added to cover the protein trap, before being incubated at 37 °C overnight in a sealed bag. Peptides were eluted into a collection tube with multiple aliquots of 40 μL increasing concentration of acetonitrile in 0.1% formic acid. Samples were spun dry and then resuspended in 20 μL buffer A (0.1% formic acid (aq)) for later analysis by mass spectrometry.

Proteomics analysis was performed using LC-MS/MS (Thermo Fisher Scientific UHPLC system coupled to an Exploris 480 mass spectrometer). The electrospray voltage was 2.2 kV in positive-ion mode. The ion transfer tube temperature was 250 °C. Full MS scans were obtained in the Orbitrap mass analyser from m/z 340 to 1110, with a mass resolution of 120,000 (at m/z 200). The automatic gain control (AGC) target value was set at ‘Standard’. The maximum accumulation time was ‘Auto’ for the MS/MS. The MS/MS ions were monitored in 12 windows from mass 350-470, in 36 windows from mass 465-645 and 10 windows from mass 640-1100. Data analysis was performed using Spectronaut against a reference proteome (UniProt ID UP000246246), and locus tags corresponded to Song *et al*. (2017, 2018).^19,20^ A Q-value cutoff value of 0.05 was applied to differential protein expressions.

Intracellular metabolome analysis was conducted based on a method developed previously for autotrophic growth of *C. autoethangenum*.^21^ 5 ODs of culture was immediately pelleted by centrifugation (16,000g for 3 minutes at 4 °C) then resuspended in chilled 50% acetonitrile to extract intracellular metabolites. Further centrifugation was used to remove cell debris and then the supernatant stored at −80 °C. Samples were freeze dried and resuspended in 90 μL 2% acetonitrile which contained a 5 μM azidothymidine standard. Lipophilic compounds (such as lipids, fatty acids, oil) were removed by adding 250 uL of chloroform, then 410 μL MQ water, and then separating, by centrifugation, the organic and polar phases. The non-polar fraction was discarded, and the polar fraction cleaned through a spin column, again freeze dried and resuspended with 2% acetonitrile. LC-MS/MS analysis was performed following Espinosa *et al*. (2020) and Mahamkali *et al*. (2020),^7,21^ with modifications and additions to the scheduled multiple reaction monitoring (sMRM) transitions (using a Shimadzu UHPLC System (Nexera X2) coupled to a Shimadu 8060 triple quadrupole (QqQ) mass spectrometer). Chromatographic separation was undertaken with a Gemini NX-C18 column (3μm x 150mm x 2mm, PN: 00F4453B0, Phenomenex) and an ion-pairing buffer system consist of mobile phase A: 7.5mM tributylamine (pH 4.95 with acetic acid) and mobile phase B: acetonitrile. Sample volumes of both 5 μL and 10 μL were injected so measured intensities were within the calibration curve.

### 2.3 Thermodynamic and kinetic metabolic flux analysis

Reaction thermodynamic driving forces were evaluated using the thermodynamic variability analysis model presented by Mahamkali *et al*. (2020), with pH assumed 1 unit higher than the extracellular pH.^14^ Measured values from intracellular metabolomics (or the lower limit of quantification, LLOQ) constrained reaction conditions. Depending on transport mechanism, carboxylate products (*i.e*., acetate, butyrate *etc*.) intracellular concentrations were limited to between *ca*. 1 times higher, or 10^6^ times lower, than extracellular concentrations.^14,21^ For the formate intracellular concentration, the equivalent was between 10 times higher, or 50 times lower, than the extracellular concentration.^23^

Minimum metabolites concentrations were set to be 0.1 μM, and 1 nM for dissolved gases unless directly measured and evaluated from Henry’s law (H_2_, CO_2_ and CO). CO_2_ total concentration, *c*, was calculated from partial pressure, *p*, as,

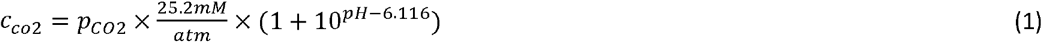

Activated metabolite maximum values were set to 1 mM, and 10 mM for others.

Reaction fluxes were estimated as follows to assess potential bottlenecks in metabolism. Metabolic flux (*J*) includes terms for protein concentration (E), Gibbs free energy of reaction (Δ*G*), saturation by substrates (*M*) with affinity (*K*), kinetic orders (*a*), and other sources of regulation (*U*), according to Heffernan *et al*. (2022),^17^

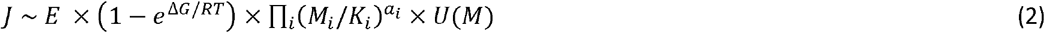

Using Michaelis-Menten enzyme kinetics, the enzyme and kinetics terms can be simplified for a given substrate concentration (*S*) to give flux (*J*) as,

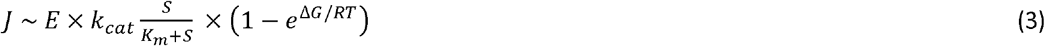

By making several further assumptions, we can get likely order of magnitude estimates for *J* as follows.

- Whilst our proteomic analysis is not able to determine absolute quantifications due to differences in MS protein constituent signal responses, *E* can still be estimated, as during a spiked protein analysis with similar setup, Valgepea *et al*. (2022),^24^ found a linear relationship between the log_10_ anchor protein concentrations and log_10_ of mass spectrometer intensity. Due to slight variability of the injected sample protein mass, we scaled protein concentrations using the pta enzyme, which was shown by Valgepea et al. (2022) to be largely constant across multiple gas fermentation conditions.
- There have been few *E. limosum* proteins assayed, and so to provide an estimate of kinetic parameters, k_cat_ and K_m_, we undertook BLAST analysis of key proteins against known protein assays. This was used to verify deep learning models predicting kinetic parameters for the WLP central carbon metabolism using protein amino acid sequences as input.^25,26^ When intracellular S was unknown, k_cat_/K_m_ was used as a proxy term for kinetic effects.

## 3 Results & Discussion

### 3.1 CO_2_ is beneficial for growth in liquid C_1_ chemostats, but detrimental for product formation

At a dilution rate (D) of 0.4 d^-1^ (specific growth rate of 0.017 h^-1^), cells reached steady-state conditions. Despite the same amount of carbon being supplied, the MeOH/CO_2_/0.4D condition (**Table 1**), reached a biomass concentration of 0.54±0.05 gDCW/L (18±3.8% of production distribution mol-C), whilst 3MeOH/Formate/CO_2_/0.4D reached a biomass concentration of 0.15±0.04 gDCW/L (7.3±1.6% mol-C) (**Table 1**). We note there was significant cell aggradation in the former condition, and so large samples were vigorously vortexed to ensure consistent biomass measurements. Methanol-specific uptake rates were higher for 3MeOH/Formate/CO_2_/0.4D – 190±23 mmol/gDCW/d compared to 75±7.0 mmol/gDCW/d for MeOH/CO_2_/0.4D (**Figure 1A**). At a comparable dilution rate, Loubiere & Lindley (1991) reached a methanol-specific uptake rate of *ca*. 72 mmol/gDCW/d (3 mmol/gDCW/h) with CO_2_ as a co-substrate. Our CO_2_-specific uptake rate was 35±14 mmol/gDCW/d (MeOH/CO_2_/0.4D), for an overall co-consumption uptake ratio of 2.6±1.4 mol-methanol/mol-CO_2_. The formate-specific uptake rate was 62±7.5 mmol/gDCW/d (3MeOH/Formate/CO_2_/0.4D), for a co-consumption uptake ratio of 3.0±0 mol-methanol/mol-formate, as expected since all liquid feedstock supplied was consumed (*i.e*., carbon limited) (**Figure 1A**). We do, however, note there was also significant CO_2_ assimilation.

**Figure 1.**
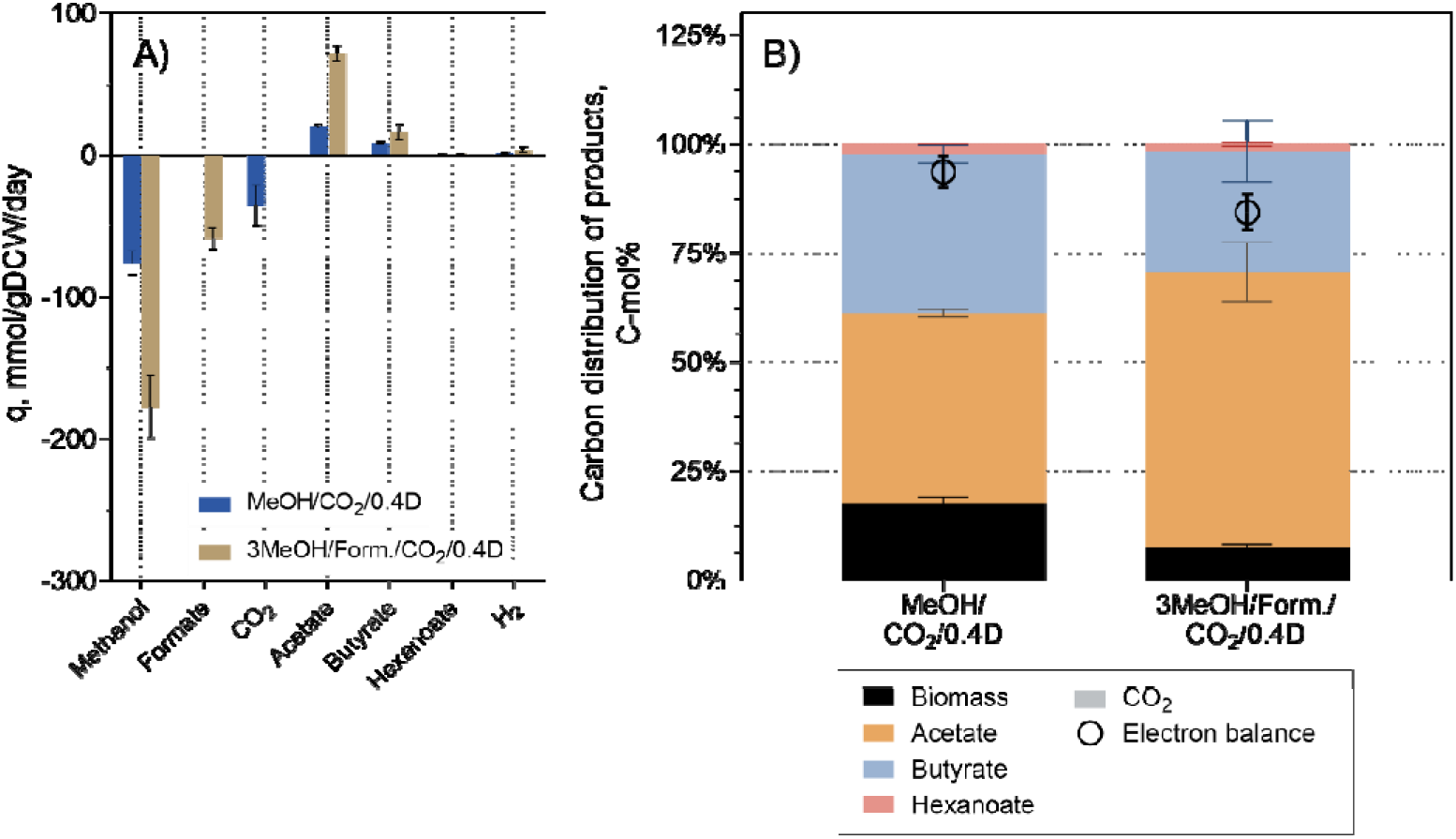
Phenomics of autotrophic (conditions as per Table 1) Eubacterium limosum chemostats. (A) specific production and uptake rates. (B) carbon distribution of products. Values are reported as the average ± standard deviation between replicates. NOTE: CO_2_ consumption has been removed for the 3MeOH/Formate/CO_2_/0.4D condition due to abiotic uncertainty, refer to text for details.

The major product of both conditions was acetate, with a specific production rate of 20±1.4 mmol/gDCW/d for MeOH/CO_2_/0.4D (43±8.7% mol-C), and 72±4.4 mmol/gDCW/d for 3MeOH/Formate/CO_2_/0.4D (63±17% mol-C) (**Figure 1A**). Butyrate was the secondary product with specific production rates of 8.5±1.0 mmol/gDCW/d (37±6.8% mol-C) and 17±4.9 mmol/gDCW/d (28±9.1% mol-C), respectively (**Figure 1A**). Hexanoate was a very minor product with production rates of 0.39±0.06 mmol/gDCW/d (2.5±0.5% mol-C) and 0.70±0.27 mmol/gDCW/d (1.5±0.5% mol-C), respectively (**Figure 1A**). Butanol was not detected in any of the tested conditions. Hydrogen was a minor sink of electrons at 1.1±0.56 mmol/gDCW/d for MeOH/CO_2_/0.4D (0.50±0.27% mol-e) and 3.7±2.3 mmol/gDCW/d for 3MeOH/Formate/CO_2_/0.4D (0.63±0.38% mol-e) (**Figure 1**).

With these production rates, we know there must be a flux of at least 47±3.5 mmol/gDCW/d acetyl-CoA for MeOH/CO_2_/0.4D and 120±12 mmol/gDCW/d acetyl-CoA for 3MeOH/Formate/CO_2_/0.4D conditions, respectively. This is important as it allows evaluation of fluxes through the methyl and carbonyl branches – it is numerically the equivalent of the flux through both branches of the WLP into the *acs* enzyme. Because the methanol uptake rate is greater than these numbers, in both conditions, we know the methyl branch of the WLP must reverse (assuming methanol enters the pathway as methyl-THF). For MeOH/CO_2_/0.4D this reverse methyl branch flux is 28±3.9 mmol/gDCW/d, whereas for 3MeOH/Formate/CO_2_/0.4D it is 68±12 mmol/gDCW/d.

Overall, this gave a carbon balance (*i.e*., substrate C recovery into products) of 93±18% for MeOH/CO_2_/0.4D. The carbon balance for 3MeOH/Formate/CO_2_/0.4D was below our acceptance threshold and therefore is not reported. Nonetheless, the electron balances were close to 100% at 94±4% and 84±4% for the two conditions, respectively (**Figure 1B**), suggesting the error is related to a carbon compound with a redox number of 0, *i.e*., CO_2_. Off-gas CO_2_ composition was analysed by two independent methods – online mass spectrometry (MS) and offline gas chromatography (GC) (see materials and methods), with good agreement between the data sets (*e.g*., for 3MeOH/Formate/CO_2_/0.4D the consumption was calculated as −46±21 mM-C/d from the GC and - 41±20 mM-C/d from the MS). Without unaccounted for electrons to assimilate CO_2_, it is reasonably sure that biotic CO_2_ consumption is significantly lower and thus is not considered further here.

### 3.2 Methanol is assimilated via methyltransferase operon

To better understand the observed phenotypes, we measured proteomics and metabolomics. We identified 1118 proteins that met the criteria for differential analysis between the MeOH/CO_2_/0.4D and 3MeOH/Formate/CO_2_/0.4D growth conditions (*i.e*. present in both samples and Q-value < 0.05). All the key enzymes of the WLP, direct acetyl-CoA condensation pathway, carboxylate/alcohol production, energy conservation, and central carbon metabolism were detected (**Figure 2, Figure 3**). 31 intracellular metabolites were identified in both conditions, however, unfortunately coenzyme A was consistently below the lower limit of quantification (LLOQ).

**Figure 2.**
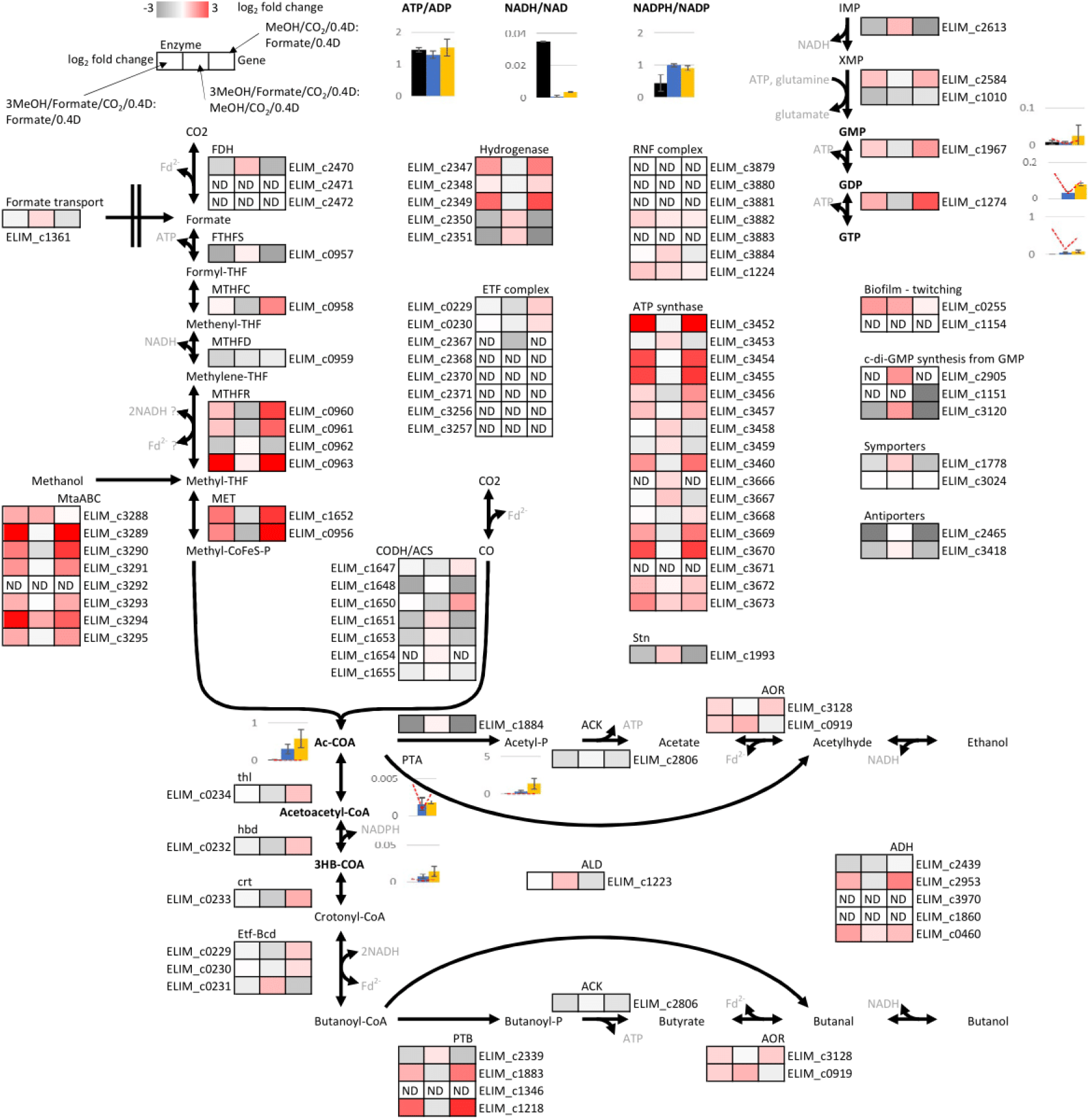
E. limosum Wood Ljungdahl Pathway metabolism schematic representation during C, fermentation. Reactions are not balanced stiochiometrically along pathways, however show substrates, products, redox mediators and proteins (elim_cxxxx). Log_2_ protein expression fold changes are calculated for the listed growth conditions. Intracellular metabolites concentrations (μmol/gDCW) are shown as the average ± standard deviation of replicates, with the LLOQ shown as a red dashed line. The Formate/0.4D,^23^ MeOH/CO_2_/0.4D, 3MeOH/Formate/CO_2_/0.4D conditions are respectively in black, blue, and yellow. Abbreviations: Ac-CoA, acetyl-CoA; ALD, aldehyde dehydrogenase; ACS, acetyl-CoA synthase; ADH, alcohol dehydrogenase; ACK, acetate kinase; AOR, aldehyde:ferredoxin oxidoreductase; Etf-Bcd, butyryl-CoA dehydrogenase; CODH, CO dehydrogenase; crt, crotonase; FDH, formate dehydrogenase; FTHFS, formyl-THF synthetase; hbd, 3-hydroxybutyryl-CoA dehydrogenase; MtaABC, methanol dependent methyl transferase; MTHFC, methenyl-THF cylcohydrolase; MTHFD, methylene-THF dehydrogenase; MTHFR, methyltransferase / methylene-THF reductase; PTA, phosphotransacetylase; PTB, phosphotransbutyrylase^19^ and therefore a butyrate kinase, BK, is assumed; Stn, Sporomusa type Nfn transhydrogenase; THF, tetrahydrofolate; thl, thiolase

**Figure 3.**
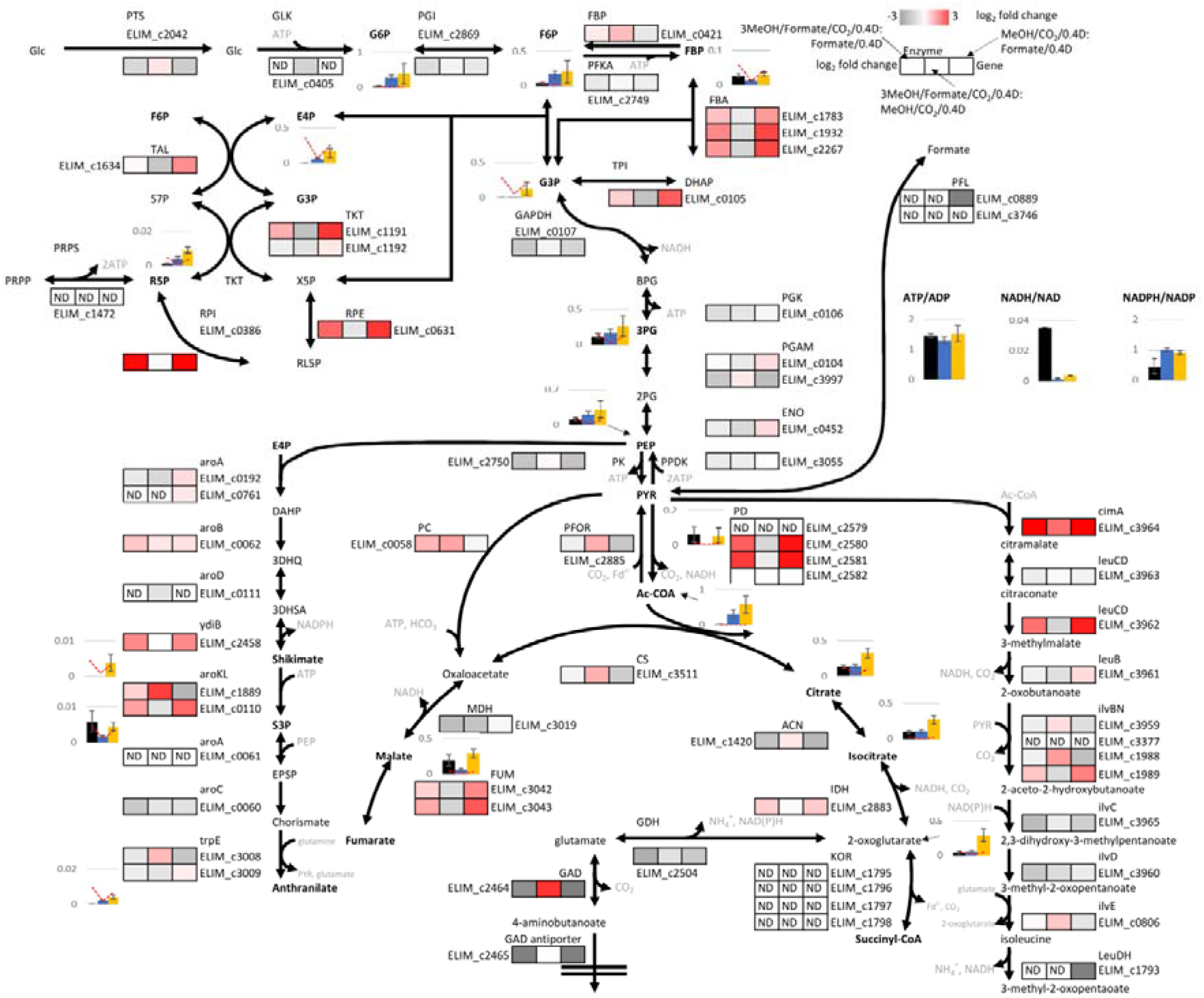
E. limosum central carbon metabolism schematic representation during C_1_ fermentation. Reactions are not balanced stiochiometrically along pathways, however show substrates, products, redox mediators and proteins (elim_cxxxx). Log_2_ protein expression fold changes are calculated for the listed growth conditions. Intracellular metabolites concentrations (μmol/gDCW) are shown as the average ± standard deviation of replicates, with the LLOQ shown as a red dashed line. The Formate/0.4D,^23^ MeOH/CO_2_/0.4D, 3MeOH/Formate/CO_2_/0.4D conditions are respectively in black, blue, and yellow. Abbreviations: 2PG, 2-phosphoglycerate; 3DHQ, 3-dehydroquinate; 3DHSA, 3-dehydroshikimate; 3PG, 3-phosphoglycerate; Ac-CoA, Acetyl-CoA; BPG, 1,3-biphosphoglycerate; DĀHP, 3-deoxy-arabino-heptulonate 7-phosphate; DHĀP, Dihydroxy acetone phosphate; E4P, Erythrose 4-phosphate; EPSP, 3-enolpyruvyl-shikimate 5-phosphate; F6P, Fructose 6-hosphate; FBP, Fructose 1,6-biphosphate; G3P, Glyceraldehyde 3-phosphate; G6P, Glucose 6-phosphate; Glc, Glucose; PEP, Phosphoenolpyruvate; PRPP, 5-phosphoribosyl diphosphate; PYR, Pyruvate; R5P, Ribose 5-phosphate; RL5P, Ribulose 5-phosphate; S3P, Shikimate 3-phosphate; S7P, Sedoheptulose 7-phosphate; X5P, Xylulose 5-phosphate; ACN, anonitase; aroA, DAHP synthase; aroA, 3-phosphoshikimate 1-carboxyvinyltransferase; aroB, 3-dehydroquinate synthase; aroC, chorismate synthase; aroD, 3-dehydroquinate dehydratase; aroKL, shikimate kinase; cimA, citramalate synthase; CS, citrate synthase; ENO, enolase; FBĀ, Fructose biphosphate aldose; FBP, fructose 1,6-biphosphatase; FUM, fumarase; GAD, glutamate decarboxylase; GAPDH, glyceraldehyde 3-phosphate dehydrogenase; GDH, glutamate dehydrogenase; GLK, Glucokinase; IDH, isocitrate dehydrogenase; ilvBN, pyruvate:2-oxobutanoate acetaldehydetransferase; ilvC, 2-aceto-2hydroxy-butanoate:NADP+ oxidoreductase; ilvD, dihydroxy acid dehydratase; ilvE, branched chain amino acid aminotransferase; KOR, oxoglutarate ferredoxin oxidoreductase; leuB, methylmalate dehydrogenase; leuCD, methylmalate hydrolase; LeuDH, leucine dehydrogenase; MDH, malate dehydrogenase; PC, pyruvate carboxylase; PD, pyruvate dehydrogenase; PFKA, Phosphofructose kinase; PFL, pyruvate formate ligase; PFOR, pyruvate ferredoxin oxidoreductase; PGAM, phosphoglycerate mutase; PGI, Glucose 6-phosphate isomerase; PGK, phosphoglycerate kinase; PK, pyruvate kinase; PPDK, phosphoenolpyruvate synthetase; PRPS, Ribose phosphat epyrophosphokinase; PTS, phosphotransferase system; RPE, Ribulose phosphate epimerase; RPI, Ribose phosphate isomerase; TAL, Transaldolase; TKT, Transketolase; TPI, Trisephosphate isomerase; trpE, anthranilate synthase; ydiB, shikimate dehydrogenase

Methanol is assimilated through the methyl transferase operon elim_c3288-3295. Compared to solely formatotrophic growth, we can see methanol assimilation results in large upregulation of most proteins in this operon, for example, elim_c3289, 3290, 3294 (log_2_ fold change 3.7, 2.3, 1.9). Elim_c3293 (mtaB), 3294 (mtaC2), 3295 (mtaA) are also the 1^st^, 3^rd^ and 10^th^ most ‘abundant’ in the 3MeOH/Formate/CO_2_/0.4D proteome. Perhaps intuitively, supplementing MeOH/CO_2_/0.4D with formate had little change on the expression of the operon – that is, regulation is controlled by the presence of methanol, not its concentration.

### 3.3 Formate triggers a beneficial intracellular state during methylotrophic growth

Supplementing MeOH/CO_2_/0.4D condition with formate produced only minor changes in the proteome, despite improving methylotrophic growth rate.^4^ This confirms that the metabolism of acetogens is not controlled at the proteomics level.

Formate offers several other useful benefits to methanol metabolism and growth rate. PFOR (elim_c2885) links the WLP carbon flux pyruvate used in glucogenesis. Adding formate to the MeOH/CO_2_/0.4D condition resulted in a slight upregulation of PFOR, however, the log_2_ fold change was <1.5. Increased PFOR activity has been noted to correlate with improved growth during a proteomics study of *E. limosum B2*.^27^ Pyruvate was not detected in the MeOH/CO_2_/0.4D condition (LLOQ 0.0016 μmol/gDCW), yet it was 0.055±0.042 μmol/gDCW in 3MeOH/Formate/CO_2_/0.4D growth. Pyruvate levels are high compared to a typical syngas fermenter, *C. ljungdahlii* in fed batch reaching *ca*. 0.02 μmol/gDCW.^28^ Interestingly, methanol metabolism appears to result in upregulation of the pyruvate-consuming PD (elim_c2580, 2581) (log_2_ fold change of 1.9 and 2.3 respectively for adding methanol/CO_2_ to formate/0.4D growth).

Formate increases pyruvate concentration due to energy limitations at the PEP level.^15,23^ This can increase flux to products downstream of pyruvate, for example, synthetic isobutanol production.^28^ Alternatively, this could also be an avenue to produce C3 compounds, of which most production pathways go through pyruvate (e.g., acrylate, methylglyoxal). Further, adding MeOH/CO_2_/0.4D to formate upregulated the propanediol dehydratase (elim_c1214, log_2_ fold change of 2.7), and is in fact, it is the 18^th^ most ‘abundant’ protein in the proteome. Hence, both formate and methanol together may be important for future synthetic C3 production.^23^

Methanol also helps overcome some of the limitations of formate metabolism. Whereas in formatotrophic growth, where the total ATP, NAD, NADP, and CoA pool sizes were low, causing stress conditions, here they are all of the same order of magnitude for MeOH/CO_2_/0.4D and 3MeOH/Formate/CO_2_/0.4D (**Figure 2, Figure 3**).^23^ For example, whilst adding formate to MeOH/CO_2_/0.4D almost doubled the acetyl-CoA pool (0.577±0.245 μmol/gDCW compared to 0.300±0.133 μmol/gDCW), adding MeOH/CO_2_ to formate/0.4D increased it more than 40-fold.

Adding formate to MeOH/CO_2_/0.4D increases FBP, acetyl-phosphate and NADH/NAD ratio, all of which are associated with solventogenesis in acetogens (**Figure 2, Figure 3**).^29^ During ethanol production, *C. autoethanogenum* had a similar acetyl-phosphate intracellular concentration to that here (1.35±0.72μmol/gDCW).^21^ The glucogenesis pathway (elim_c3055, 0452, 3997, 0104, 0106, 0107, 0105, 1783, 1932, 2267, 0421, 2869) all have log_2_ fold changes less than 1.5. Two notable exceptions are RPI (elim_c0386) and RPE (elim_0631) in the pentose phosphate pathway, which are upregulated when adding MeOH/CO_2_ to formate/0.4D (log_2_ *ca*. 2-3). Of all the measured intermediates, all were higher in the 3MeOH/Formate/CO_2_/0.4D condition (compared to MeOH/CO_2_/0.4D) – at least 2x higher in the pentose phosphate pathway (E4P, R5P, G3P), and 1-1.5x higher in glucogenesis (G6P, F6P, 3PG, PEP, except for FBP). Together these results suggest that the combined 3MeOH/Formate/CO_2_/0.4D metabolism triggers increased flux through the pentose phosphate pathway, which are known precursors for histidine, a component of corrinoid binding proteins.^12^ It is possible this increases the methanol assimilation capacity, as well as growth rate. This all suggests a small amount of formate is needed for solventogenesis, and improved methanol uptake.

### 3.4 Butyrate production feasible only if hbd is NADPH dependant

There is mostly little change in expression of proteins that can produce carboxylates from acetyl-CoA (elim_c0231-0234, 2806, 2339); elim_c0233 is the most downregulated with a log_2_ fold change of 1.2 between 3MeOH/Formate/CO_2_/0.4D and MeOH/CO_2_/0.4D. Acetoacetyl-CoA and 3-hydroxybutyrate-CoA had similar intracellular concentrations (0.672±0.081μmol/gDCW versus 0.594±0.316μmol/gDCW and 5.24±2.52μmol/gDCW versus 2.83±1.04μmol/gDCW respectively). Since there was a difference in butyrate production between the conditions, this reinforces that metabolism is regulated post-translation. In both conditions, the measured concentrations of acetoacetyl-CoA, 3-hydroxybutanoyl-CoA, and NADH/NAD mean hbd is not thermodynamically feasible for butyrate production. However, since this was observed, we suggest hbd is NADP-dependant, which we measured to be far more reduced than NAD (**Figure 2**). **Figure 4** shows this NADP-dependant hbd has sufficient thermodynamic driving force to make butyrate production feasible.

**Figure 4.**
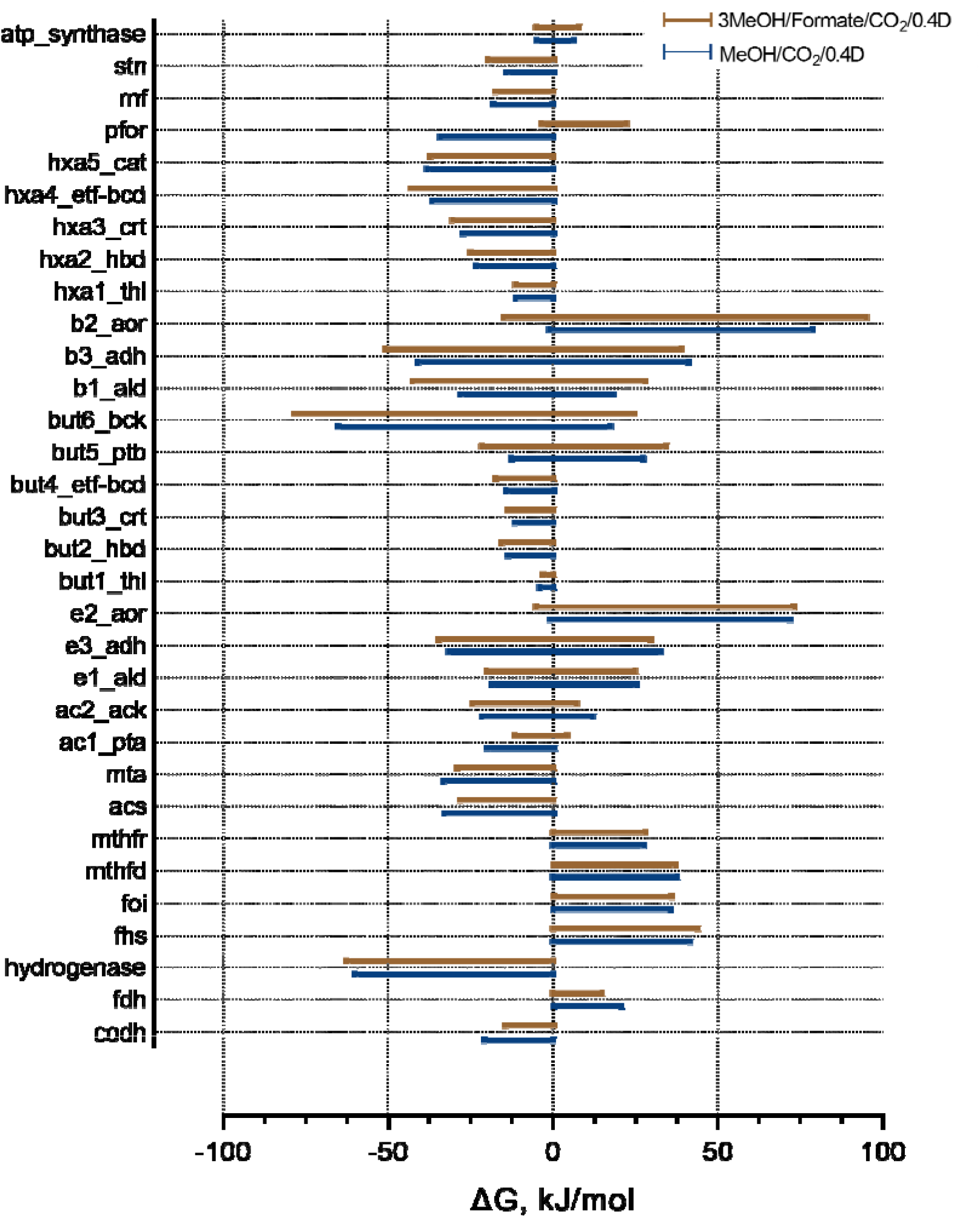
E. limosum thermodynamic variability analysis showing maximum allowable Gibbs free energy range for metabolic reactions. MeOH/CO_2_/0.4D is shown in blue, and 3MeOH/Formate/CO_2_/0.4D in brown. Reactions are calculated in the anabolic direction from CO_2_ shown on Figure 2, Figure 3. Rnf, stn, hydrogenase and PFOR are calculated for the ferredoxin-consuming direction. ATP synthase is calculated for the ATP-forming direction.

### 3.5 Rnf direction uncertainty

During autotrophic acetogen growth, including formate, membrane bound Rnf proteins couple oxidation of reduced ferredoxin to the reduction of NAD to generate a transmembrane ion gradient for ATP synthesis (*i.e*. chemiosmotic transport). However, there is a lack of reduced ferredoxin on low energy substrates, such as ethanol and methanol, which only generate NADH during oxidation. Thus, authors have speculated that Rnf may operate in the reverse direction,^12^ with the transmembrane ion gradient established by ATP hydrolysis. Therefore, one might expect a difference in Rnf expression for methanol and formate conditions (due to the supposed opposite function of Rnf), which we did not observe (**Figure 2**). However, since Rnf has activity orders of magnitude lower than ATPase,^30^ and activity does not correlate with expression,^31^ proteomics data alone is not sufficient to refute the hypothesis.

Rnf and ATPase from acetogens have been purified into liposomes, where ATP hydrolysis was able to generate large transmembrane ion gradients for endergonic reduction of ferredoxin (*i.e*. operating in the reverse direction).^30^ We however note these conditions – pH 7.5, with the reaction ceasing at NADH/NAD = 110, and ATP/ADP = 88, are very different to physiological conditions measured here (*e.g*. respectively at 7.8 intracellular, 0.0036, 1.5 respectively for 3MeOH/Formate/CO_2_/0.4D). So, whilst Rnf can theoretically reverse, which would help satisfy the reduced ferredoxin demand for methylotrophic metabolism, it is not thermodynamically feasible under the conditions tested (**Figure 4**). There are limited conditions where it might occur, for example, at high pH, or during non-steady state growth when the NADH pool is more reduced, such as seen in early syngas growth phases by Mahamkali and colleagues.^21^ As a result of this thermodynamic constraint, what we think we know about *E. limosum* metabolism must be revisited, particularly around this energy conservation.

### 3.6 Energy conservation

There was little change in bifurcating-etf proteins (elim_c0229, 0230, 2367), nor Rnf proteins (elim_c3882, 3884). ATP/ADP intracellular ratios were comparatively consistent at 1.5±0.26 for 3MeOH/Formate/CO_2_/0.4D, 1.3±0.11 for MeOH/CO_2_/0.4D, as were the NADPH/NADP (0.91±0.072 and 1.0±0.056) and NADH/NAD ratios (0.0036±0.0004 and 0.0011±0.0006). The sporomusa type Nfn complex (elim_c1993)^32^ was upregulated, but insignificantly, when formate was added to MeOH/CO_2_/0.4D with a log_2_ fold change of <1.5. The total ATP, NAD, NADP, and CoA pool sizes are all of same order of magnitude for MeOH/CO_2_/0.4D and 3MeOH/Formate/CO_2_/0.4D, although generally slightly larger for the latter. The total FMN pool (including FAD) is approximately 5% of a typical acetogen growing on syngas.^21^

We could not directly measure ferredoxin, however since ferredoxin is used in several reactions, we could define lower and upper bounds, here discussed for thermodynamics of the 3MeOH/Formate/CO_2_/0.4D condition. From the Stn, Fd^2-^/Fd must be > 0.4, assuming there is a downstream significant use of NADPH. Dissolved hydrogen, calculated using Henry’s law, is *ca*. 0.26 μM, which hyt suggests Fd^2-^/Fd must be > 3. With the Rnf/ATPase complex working in the reverse direction for methanol metabolism (*i.e*. consuming ATP to produce ferredoxin required for methylene-tetrahydrofolate reductase (MTHFR), assuming it is bifurcating), Fd^2-^/Fd should be < 0.4. For PFOR to be generating pyruvate, Fd^2-^/Fd must be >1. Taken together, this implies an apparent conflict in the required ferredoxin concentration. In our view, this confirms Rnf that can not reverse, and so is not a source of reduced ferredoxin in methylotrophic growth.

Unlike Rnf, we found that MTHFR was regulated with ATPase (**Figure 2**), and so perhaps MTHFR is also involved in chemiosmotic transport. This has previously been postulated by researchers, as the reduction of methylene-THF to methyl-THF has a more positive redox potential than NADH (−200mV versus −320mV, respectively).^33^ However, evidence of how energy is conserved, whether by bifurcation or chemiosmotic electron transport, is still missing.^34^ Hence the MTHFR is considered one of the biggest uncertainties in metabolism and has implications for ferredoxin balance.^12^

Since we know the fluxes through ferredoxin-consuming pathways (CODH, hyt, PFOR for biomass) and ferredoxin production pathways (fdh, etf-Bcd), we can calculate the experimental ferredoxin production that is available for chemiosmotic ATP generation. For simplicity, we have not included MTHFR in this calculation. Using current knowledge of *E. limosum* metabolism,^4^ the Fd^2-^/Fd is likely *ca*. >3 for both MeOH/CO_2_/0.4D and 3MeOH/Formate/CO_2_/0.4D conditions, which will cause a flux through the Stn, leading to NADPH. Given we found hbd must be NADPH dependent, this does make sense and perhaps explains why there is a relatively reduced NADP pool, especially compared to typical syngas fermentations.^21^ Most importantly, this extra NADPH demand means there is an extra ferredoxin demand to account for (consumed by Stn). Consequently, for MeOH/CO_2_/0.4D there is a net ferredoxin requirement of 24±2.5 mmol/gDCW/d, and for 3MeOH/Formate/CO_2_/0.4D a production of 5.5±9.9 mmol/gDCW/d. Thus, it is clear that a separate source of ferredoxin is required, or alternatively, demand is overestimated. In either case, this data likely suggests sufficient reduced ferredoxin is not available for a bifurcating-MTHFR during methylotrophic growth.

### 3.7 Methylene reductase: the missing link in energy conservation?

*E. limosum* contains a type II MTHFR, which comprises a MetF (elim_c0960; downregulated log_2_ fold change of 1.5 when adding formate to MeOH/CO_2_/0.4D) with a ferredoxin binding site MetV (elim_c0961). In the direction producing methyl-THF (*i.e*. as in formate metabolism), this would be very energetically wasteful, whilst in the reverse direction (*i.e*. for methanol metabolism), the energetic barrier would make the WLP impossible. Öppinger et al (2022) proposed a potential ‘bifurcating’ solution to this using the adjacent, and highly expressed LpdA and GcvH, which currently have unknown functions.^24,34^ Perhaps the upregulated GcvH is the electron acceptor for methyl-THF oxidation, given it is significantly upregulated when adding methanol/CO_2_ to formate/0.4D (*i.e*., when the methyl branch must reverse) (elim_c0963; 11^th^ most upregulated protein, log_2_ fold change 4.9). Öppinger et al (2022) proposed this was coupled with NAD via LpdA (elim_c0962) as the terminal acceptor, yet we found LpdA was downregulated when adding methanol/CO_2_ to formate/0.4D (log_2_ fold change of −1.2), thus conclude this is unlikely how MTHFR oxidises methyl-THF.

Alternatively, it has been proposed for MTHFR to electrically connect to a Rnf. MTHFR is generally in the cytoplasm, although has been observed in the membrane fraction.^35^ It is unclear how this applies to *E. limosum*, yet, thermodynamically, there is no reason this would not work with our observed data. Perhaps most interestingly, both MeOH/CO_2_/0.4D and 3MeOH/Formate/CO_2_/0.4D can achieve overall ferredoxin balance without requiring different metabolisms. This electrically connected MTHFR-Rnf does, however, require methyl-THF intracellular concentration to be approximately four orders of magnitude higher than methylene-THF and methenyl-THF. Measurement of these intracellular metabolites is difficult, and to our knowledge, has only been done once, for syngas fermentation,^36^ yet may provide further clues as to the MTHFR function in methylotrophic growth.

### 3.8 Formate and high dilution rate increase methylotrophic productivity

Since we detected more acetate relative to butyrate here than in our previous batch experiments,^4^ the methanol-to-formate uptake ratio was not high enough to divert flux to more reduced products. However, we know that the methanol uptake rate increases more relative to other substrates as the dilution rate increases. Therefore, we conducted further experiments by increasing the methanol feed to a methanol:formate ratio of 10:1 at a dilution rate of 1 d^-1^ (10MeOH/Formate/CO_2_/1D). With the steady-state conditions thus far, CO_2_ played a much bigger role than seen in batch and was detrimental to reduced product formation. Therefore, we aimed to understand if CO_2_ could be removed as another means to improve flux to butyrate (10MeOH/Formate/1D). We also evaluated the effect of increasing the dilution rate further to 2 d^-1^ (10MeOH/Formate/2D).

Cells reached steady-state conditions at a dilution rate of 1.0 d^-1^ (specific growth rate of 0.042 h^-1^), 10MeOH/Formate/CO_2_/1D, with a biomass concentration of 0.20±0.04 gDCW/L (12±6.3% of production distribution mol-C) (**Table 1**). We then removed the CO_2_ gas feed at the same dilution rate (10MeOH/Formate/1D), with biomass eventually stabilising at 0.10±0.01 gDCW/L (8.0±2.9% mol-C) (**Table 1**). However, we highlight that achieving growth without CO_2_ was difficult, suggesting the strain is not well adapted to this phenotype. We found the most successful strategy was to grow cells on methanol and CO_2_, then slowly remove CO_2_ as formate was added. This proves CO_2_ is not required to balance the culture metabolism. However, it clearly has a significant benefit in terms of biomass production.

Given our previous batch experiments showed formate as a co-substrate had advantages over CO_2_, we investigated whether a methanol and formate culture could reach higher dilution rates. We achieved a steady state at 2 d^-1^, 10MeOH/Formate/2D, surprisingly finding this increased the biomass concentration to 0.13±0.04 gDCW/L (8.7±1.3% mol-C) (**Table 1**). Methanol specific uptake rates were respectively 250±70 mmol/gDCW/d, 330±150 mmol/gDCW/d and 640±22 mmol/gDCW/d for 10MeOH/Formate/CO_2_/1D, 10MeOH/Formate/1D and 10MeOH/Formate /2D (**Figure 5**). This compares to a maximum for Loubiere & Lindley (1991) of 330 mmol/gDCW/d at washout for methanol/CO_2_ growth.^16^ A mixotrophic growth of methanol/glucose had a maximum methanol uptake of 120 mmol/gDCW/d at a dilution rate of 3.6 d^-1^.^37^ These results suggest that adding formate as a methanol co-substrate significantly improves methanol uptake, which is in line with our observations.

**Figure 5.**
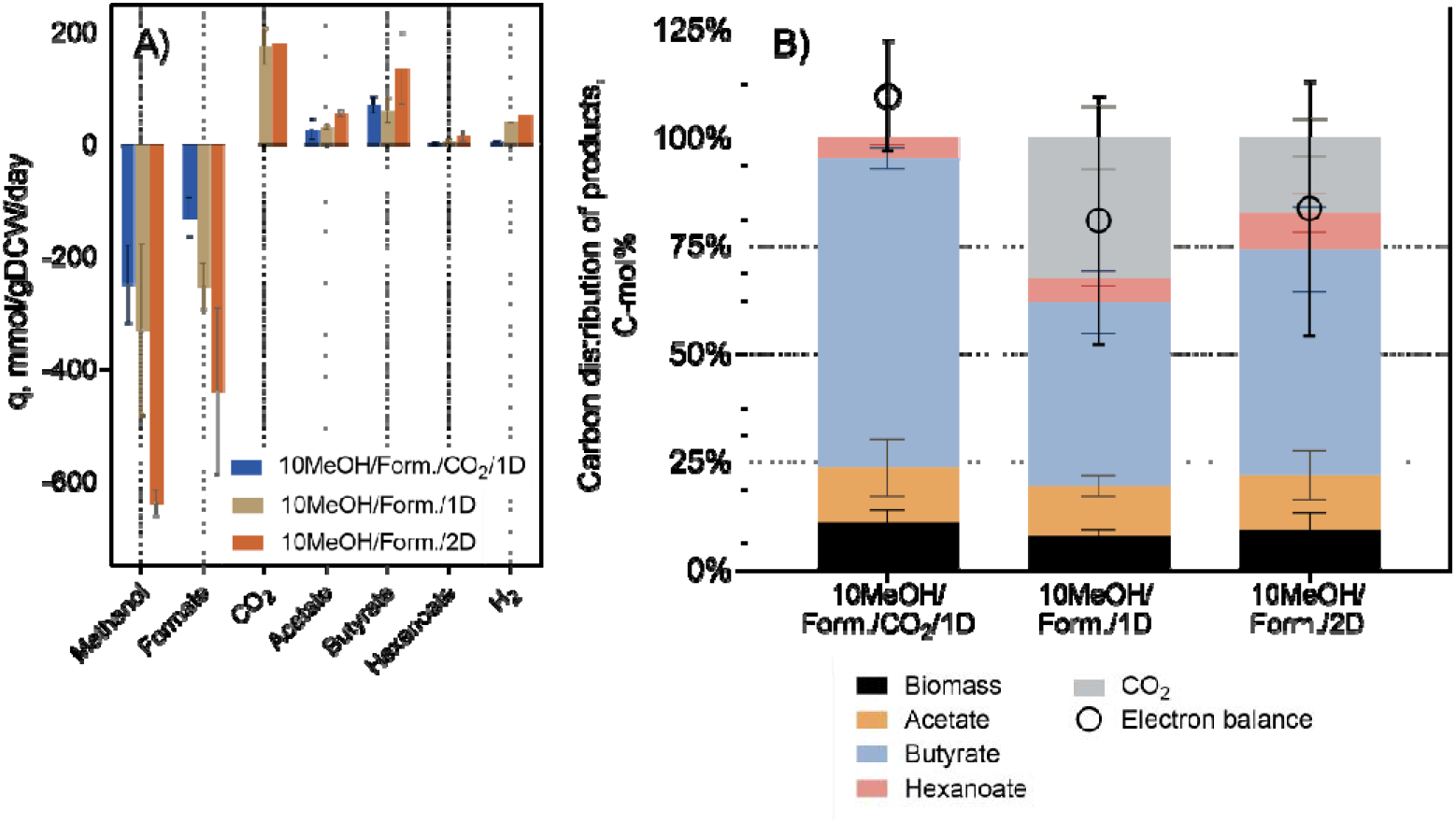
Phenomics of autotrophic (conditions as per Table 1) Eubacterium limosum chemostats. (A) specific production and uptake rates. (B) carbon distribution of products. Values are reported as the average ± standard deviation between replicates. NOTE: CO_2_ consumption is not shown for the 10MeOH/Formate/CO_2_/1D condition due to abiotic uncertainty, refer to text for details.

Somewhat intriguingly, the methanol-to-formate co-consumption uptake ratio was lower than the previous set of experiments when the methanol feed was increased (1.9±0.3 for 10MeOH/Formate/CO_2_/1D versus 3.0±0 mol-methanol/mol-formate for 3MeOH/Formate/CO_2_/0.4D). However, this may also be related to a difference in pH setpoint (7.3 versus 6.8). As expected, removing CO_2_ from the gas feed (10MeOH/Formate/1D) further reduced the co-consumption (mol-methanol/mol-formate) uptake rate to 1.3±0.5, while increasing the dilution rate (10MeOH/Formate/2D) increased it to 1.6±0.7, confirming the importance of a high dilution rate for methylotrophy. Interestingly, methanol is also beneficial for formate metabolism – the highest formate specific uptake rate of 440±150 mmol/gDCW/d (**Figure 5A**) is higher than during formate-only metabolism as a sole substrate.^23^ Together these results show we achieved a maximum electron uptake rate of 55±5 μmol/gDCW/s, which is higher than observed in CO metabolism (*ca*. 4 – 25 μmol/gDCW/s), and comparable to the mean of hydrogen uptake (*ca*. 50 μmol/gDCW/s).^38^

Unlike before, the major product in these three conditions was butyrate, with specific production rates of 70±15 mmol/gDCW/d for 10MeOH/Formate/CO_2_/1D (72±26% mol-C), 59±22 mmol/gDCW/d for 10MeOH/Formate/1D (42±7.3% mol-C), and 130±60 mmol/gDCW/d for 10MeOH/Formate/2D (54±20% mol-C) (**Figure 5A**). Not surprisingly, our maximum butyrate production was more than twice that achieved in previous batch fermentations.^4^ Acetate was the secondary product with butyrate:acetate ratios (mol-C basis) of 7.3±3.9, 3.8±1.1 and 5.0±2.5, respectively. When CO_2_ was removed as a substrate, there was significant production of 170±33 mmol/gDCW/d at a dilution of 1 d^-1^ (33±11% mol-C) (10MeOH/Formate/1D) and 180 mmol/gDCW/d at 2 d^-1^ (17% mol-C) (10MeOH/Formate/2D) (**Figure 5A**). This indicates that perhaps there is a minimum intracellular CO_2_ concentration required for metabolism, or an imbalance between production/uptake kinetics.

From the observed production rates, we know the methyl branch of the WLP must reverse (assuming methanol enters the pathway as methyl-THF). For 10MeOH/Formate/CO_2_/1D the flux is 50±38 mmol/gDCW/d, for 10MeOH/Formate/1D the flux is 140±110 mmol/gDCW/d, whereas for 10MeOH/Formate/2D it is 230±150 mmol/gDCW/d. Compared to the previous set of experiments, these highlight that removing CO_2_ and increasing the dilution rate are useful strategies for increasing flux through the reverse WLP methyl branch, an important characteristic for formation of reduced products.

In general, it appears methanol goes to butyrate, with excess electrons to hydrogen, and formate goes to acetate and CO_2_. Electron balances were 110±13%, 81±29%, and 83±30% for 10MeOH/Formate/CO_2_/1D, 10MeOH/Formate/1D and 10MeOH/Formate/2D, respectively (**Figure 5A**). The abiotic CO_2_ issues discussed previously again applied only to 10MeOH/Formate/CO_2_/1D growth (*i.e*., when CO_2_ was supplied as a co-substrate along with formate), with carbon balances for the latter two conditions of 98±21% and 80±26%.

### 3.9 Barriers to butanol formation

Unlike butyrate, butanol has a sizeable chemical market, and holds promise for use as a drop-in fuel.^11^ However, butanol was below the LLOQ in all cultures here. From a stoichiometric perspective, butanol formation requires a very high methanol:formate consumption ratio, higher than butyrate. However, when that was tested in chemostat here, it appears the extra reducing equivalents were instead directed to hexanoate and in some cases, hydrogen. We observed production hexanoate rates of 3.0±0.33 mmol/gDCW/d (resulting in a carbon distribution of 5.2±3.2% mol-C), 5.1±2.7 mmol/gDCW/d (5.4±1.6% mol-C) and 15±11 mmol/gDCW/d (9.0±5.7% mol-C) for 10MeOH/Formate/CO_2_/1D, 10MeOH/Formate/1D and 10MeOH/Formate/2D respectively (**Figure 5A** and **B**). When CO_2_ was provided as a substrate (10MeOH/Formate/CO_2_/1D), there was little flux to hydrogen as an electron sink (4.3±0.7 mmol/gDCW/d; 0.40±0.23% mol-e). However, when CO_2_ was removed, flux to hydrogen increased more than 10-fold to 39±1.6 mmol/gDCW/d (3.7±2.0% mol-e) (10MeOH/Formate/1D) and 51 mmol/gDCW/d (2.0% mol-e) (10MeOH/Formate/2D) (**Figure 5A** and **B**).

In batch, a 3:1 methanol:formate substrate ratio achieved a butyrate:acetate (mol-C basis) production ratio of 2.8±0.13 averaged over two growth phases (methanol and formate co-consumption, then methanol only).^4^ In chemostat we could replicate this average methanol:formate ratio only when CO_2_ was provided, indicating it is required to balance metabolism. Yet in batch during this first growth phase, there was a maximum co-consumption uptake ratio of 1:1, which was similar to the maximum achieved in chemostat when CO_2_ was removed – 1.6±0.7:1 with a butyrate:acetate production ratio of 5.0±2.5 and no butanol observed (10MeOH/Formate/2D) (**Figure 5**). In the second phase of growth in batch, when formate was exhausted, the butyrate:acetate production ratio was 17±2.7, with a butanol titer of 0.8 mM-C.^4^ In chemostat, we could not mimic this second phase of growth, highlighting this is likely an overflow metabolism.

We suspect this lower amount of butyrate in continuous is because pta expression and acetyl-coa concentration control the acetate/butyrate ratio. In MeOH/CO_2_/0.4D growth, the acetyl-CoA/CoA ratio is more than twice that for the 3MeOH/Formate/CO_2_/0.4D condition (11±5.4 versus 4.6±5.1), reflecting a larger thermodynamic driving force (note this assumes the LLOQ for CoA; **Figure 4**). Conversion to the carboxylate is catalysed by pta/ptb (elim_c1218, 1883, 1884, 2339). Since we found little change in expression of pta and ak, even those known to be more specific for butyrate,^14^ some flux will always get lost to acetyl-phosphate (0.294±0.177 μmol/gDCW for MeOH/CO_2_/0.4D versus 1.35±0.72 μmol/gDCW 3MeOH/Formate/CO_2_/0.4D) and then acetate, and this relates to thermodynamic driving force. Hence, this is why acetate production was lower for MeOH/CO_2_/0.4D than 3MeOH/Formate/CO_2_/0.4D.

In batch, almost exclusive butyrate production is achieved when formate is exhausted because of degradation of the cell energy status and slowing of growth rate.^15^ This can not be replicated in chemostat, but perhaps it could be in retentiostat. In fact, significant butanol production (from CO) with closely related *Butyribacterium methylotrophicum* was only noted in retentiostat growth.^39^ Alternatively in that same organism, overexpressing the butyrate production pathway relative to pta increased the butyrate:acetate production ratio.^11^ In this way, it may actually be more likely the methanol:co-substrate uptake ratio is partially controlled by the butyrate:acetate production ratio, not completely the other way around.

In syngas fermentations, alcohols are predominately formed via the aldehyde:ferredoxin oxidoreductase (*AOR*) enzyme.^29^ *AOR* activity is essential to consume reduced ferredoxin and maintain redox balance to control metabolic homeostasis.^21^ As we have discussed, methylotrophic fermentation has a shorter supply of reduced ferredoxin, and thus there is less driver for AOR to promote alcohol formation. It is also likely the ferredoxin pool is less reduced here, because in syngas fermentation ferredoxin is kept reduced by codh activity. Further, methylotrophic fermentations have always been at higher pH’s than required to promote this AOR capability (>6). It is unclear whether this growth requirement is related to methanol metabolism in the WLP, or is species specific. From our model, we see no thermodynamic reason why methanol uptake is not possible at pH < 6, albeit depending on MTHFR assumption (e.g., for a MTHFR coupled with Rnf). Other reasons could prevent methylotrophic growth at low pH, such as enzyme activity, kinetics, and inhibition.^34^ We note that there could be a significant loss of electrons to hydrogen as hyt becomes more favourable at low pH, and to prevent its formation, the NAD pool would need to be more oxidised. While we did observe hydrogen production, other researchers have suggested hydrogen production capacity is not always present.^16^

### 3.10 Enzyme kinetics and thermodynamic driving forces prevent the production of butanol

Even though there was no alcohol production in any condition, key proteins were still detected including an acetylating aldehyde dehydrogenase (elim_c1223), AOR (elim_c3128, 0919) and alcohol dehydrogenase (elim_c2439, 2953, 0460). There was however little regulation between conditions, (log2 fold change < 1.5), although AOR (elim_c3128) was in the top 5% of proteome ‘abundance’. Unlike *C. ljungdahlii*, a typical syngas fermenter, our *E. limosum* proteome showed a much higher ‘abundance’ of ald relative to pta (0.3 compared to 0.01).^29^ Whereas syngas fermentations have intense repression of ald,^21^ this is not the case for *E. limosum*. This may initially seem promising because the pathway via ald is more feasible at our experimental pH (**Figure 4**). Yet, this relatively high expression of both adh and AOR in the *E. limosum* proteome causes a potential conflict. Here, ald can consume acetyl-CoA and butyryl-CoA to produce aldehyde intermediates, and in fact can be much more thermodynamically feasible than the competing pta reaction towards carboxylates (**Figure 4**). But then with flux going via ald, there is further competition by adh to produce alcohols, or AOR to run in reverse and produce carboxylates (**Figure 4**). Therefore, it becomes a question of kinetics as to where the carbon flux actually goes.

**Table 2** presents a review of published kinetic parameters for proteins with the highest identity to *E. limosum*. The ratio of k_cat_/K_m_ is a proxy for substrate conversion speed which we used to compare to the deep learning prediction models. As can be seen, this showed promising order of magnitude correlation, with few exceptions. Importantly, it highlights AOR is likely faster than ald, and thus buildup of the butanal intermediate is facilitated by AOR, not ald. However because the latter has a more negative thermodynamic driving force (**Figure 4**), a point will be reached in batch when the AOR driving force approaches zero, and the butanal concentration is maintained by ald. A flux overflow could be directed to butanol, depending on the NADH/NAD ratio becoming more reduced than measured here. This perhaps is the case in batch as observed by Lebloas and colleagues (1996) (0.088 compared to 0.0011±0.0006 in chemostat here). Kinetically, butanol production would however be limited by the relatively low k_ca_t/K_m_ values of the native ald and adh in *E. limosum* compared to the rest of metabolism **(Table 2).**

**Table 2.**
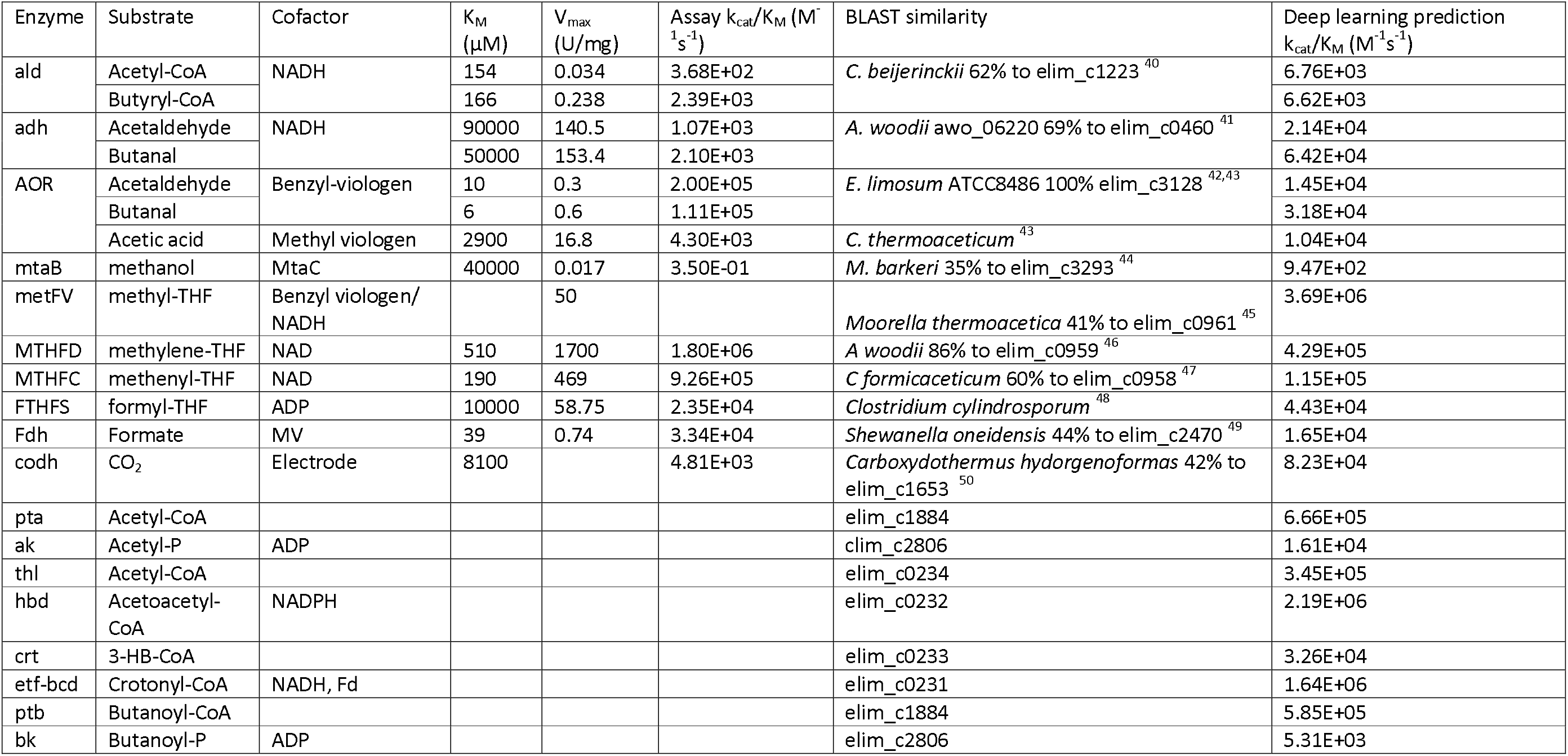
Kinetic parameters of assayed enzymes with highest BLAST similarity to E. limosum, to our knowledge, compared to deep learning prediction models.

| Enzyme | Substrate | Cofactor | K<sub>M</sub><br>(μM) | V<sub>max</sub><br>(U/mg) | Assay k<sub>cat</sub>/K<sub>M</sub> (M<sup>-1</sup>s<sup>-1</sup>) | BLAST similarity | Deep learning prediction<br>k<sub>cat</sub>/K<sub>M</sub> (M<sup>-1</sup>s<sup>-1</sup>) |
| --- | --- | --- | --- | --- | --- | --- | --- |
| ald | Acetyl-CoA | NADH | 154 | 0.034 | 3.68E+02 | *C. beijerinckii* 62% to elim_c1223<sup>40</sup> | 6.76E+03 |
|  | Butyryl-CoA |  | 166 | 0.238 | 2.39E+03 |  | 6.62E+03 |
| adh | Acetaldehyde | NADH | 90000 | 140.5 | 1.07E+03 | *A. woodii* awo_06220 69% to elim_c0460<sup>41</sup> | 2.14E+04 |
|  | Butanal |  | 50000 | 153.4 | 2.10E+03 |  | 6.42E+04 |
| AOR | Acetaldehyde | Benzyl-viologen | 10 | 0.3 | 2.00E+05 | *E. limosum* ATCC8486 100% elim_c3128<sup>42,43</sup> | 1.45E+04 |
|  | Butanal |  | 6 | 0.6 | 1.11E+05 |  | 3.18E+04 |
|  | Acetic acid | Methyl viologen | 2900 | 16.8 | 4.30E+03 | *C. thermoaceticum*<sup>43</sup> | 1.04E+04 |
| mtaB | methanol | MtaC | 40000 | 0.017 | 3.50E-01 | *M. barkeri* 35% to elim_c3293<sup>44</sup> | 9.47E+02 |
| metFV | methyl-THF | Benzyl viologen/<br>NADH |  | 50 |  | *Moorella thermoacetica* 41% to elim_c0961<sup>45</sup> | 3.69E+06 |
| MTHFD | methylene-THF | NAD | 510 | 1700 | 1.80E+06 | *A. woodii* 86% to elim_c0959<sup>46</sup> | 4.29E+05 |
| MTHFC | methenyl-THF | NAD | 190 | 469 | 9.26E+05 | *C. formicaceticum* 60% to elim_c0958<sup>47</sup> | 1.15E+05 |
| FTHS | formyl-THF | ADP | 10000 | 58.75 | 2.35E+04 | *Clostridium cylindrosporum*<sup>48</sup> | 4.43E+04 |
| Fdh | Formate | MV | 39 | 0.74 | 3.34E+04 | *Shewanella oneidensis* 44% to elim_c2470<sup>49</sup> | 1.65E+04 |
| codh | CO<sub>2</sub> | Electrode | 8100 |  | 4.81E+03 | *Carboxydotherrmus hydorgenofomas* 42% to elim_c1653<sup>50</sup> | 8.23E+04 |
| pta | Acetyl-CoA |  |  |  |  | elim_c1884 | 6.66E+05 |
| ak | Acetyl-P | ADP |  |  |  | elim_c2806 | 1.61E+04 |
| thl | Acetyl-CoA |  |  |  |  | elim_c0234 | 3.45E+05 |
| hbd | Acetoacetyl-CoA | NADPH |  |  |  | elim_c0232 | 2.19E+06 |
| crt | 3-HB-CoA |  |  |  |  | elim_c0233 | 3.26E+04 |
| etf-bcd | Crotonyl-CoA | NADH, Fd |  |  |  | elim_c0231 | 1.64E+06 |
| ptb | Butanoyl-CoA |  |  |  |  | elim_c1884 | 5.85E+05 |
| bk | Butanoyl-P | ADP |  |  |  | elim_c2806 | 5.31E+03 |

## 4 Conclusions

We have obtained a baseline methylotrophic phenomics, proteomics and metabolomics steady-state dataset for *E. limosum*. Formate, as opposed to CO_2_, proved to be a better co-substrate, allowing higher growth rates and methanol assimilation. In addition, a small amount of formate increased parameters such as NADH/NAD during methylotrophic growth, which correlate with solventogenesis in acetogens. Despite these positives, a bottleneck in metabolism appears to prevent further increases in methanol uptake required for alcohol production, including loss of acetyl-CoA to acetate. More importantly, kinetics and thermodynamics of AOR, ald and adh prevent production of butanol.

## Data availability

Data available upon request from the authors.

## Acknowledgements

JW acknowledges support of the Warwick and Nancy Olsen Scholarship. BV acknowledges the support of the Australian Research Council (ARC) through grants FL170100086 and LP200200136. EM acknowledges the support of the ARC CoE in Synthetic Biology CE200100029 and LP200200136.

The research utilised equipment and support provided by the BPA-funded facility, Queensland Metabolomics And Proteomics (Q-MAP), an Australian Government initiative being conducted as part of the National Collaborative Research Infrastructure Strategy (NCRIS) National Research Infrastructure for Australia.

